# The Kinase-Dependent and Independent Functions of Cdk4 and Cdk6 Regulate Continuous Proliferation and the Exit From Quiescence Differently

**DOI:** 10.1101/2020.04.23.058347

**Authors:** Sarah E. Nataraj, Stacy W. Blain

## Abstract

Cdk4 and cdk6 have long been considered functional homologues, despite the fact that tissue-specific differences are detected in the single cdk4 or cdk6 knockout animals. To explore the role of cdk4 and cdk6 in the same model system, we overexpressed variants of cdk4 and cdk6 in Mv1Lu cells to determine their effect on cell cycle progression. We found that cells that overexpressed cdk4 were able to reenter the cell cycle from a contact arrested quiescent state in a kinase-dependent fashion, consistent with the role of this kinase in G0-G1 phase exit. However, we also found that expression of catalytically inactive variants of cdk6 accelerated G0 release, enabled the cell to overcome TGF-β-mediated growth arrest, proliferate in the absence of cyclin D-associated kinase activity and maintain cdk2 activity, while cells expressing catalytically inactive cdk4 were not. This suggested that cdk6 expression was able to affect cell cycle progression in a kinase-independent manner that was distinct from the actions of cdk4. Cdk4 appears unable to associate with cyclin D in the absence of a required assembly factor, such as p27Kip1, but, by gel filtration chromatography, we detected the presence of a previously unidentified, catalytically inactive cyclin D-cdk6 dimer, which might contribute to the kinase-independent, pro-proliferative activity of cdk6.

---

The mammalian cell cycle is an ordered process where the entire genome of a cell is duplicated and then distributed equally between two daughter cells. The progression between cell cycle phases is regulated by the cyclin-cdks, where cyclin D-associated cdk4 or cdk6 starts the cycle, causing release from G0 phase (1). The presence of mitogens leads to the synthesis of the D-type cyclins, which partner with either cdk4 or cdk6 (2). The cyclin D-cdk4 or cyclin D-cdk6 complexes phosphorylate substrates, like the retinoblastoma (Rb) protein, which are involved in progression through the G1 phase (3). Antiproliferative responses, such as growth to confluence or TGF-β treatment, cause inhibition of cyclin D-cdk4 and cyclin D-cdk6 complexes by the association of cyclin-dependent kinase inhibitors (CKIs), such as p15^Ink4b^, p27^Kip1^, or p21^Cip1^ (4, 5).

In addition to its catalytic function, cyclin D-cdk4/6 complexes have at least one kinase-independent function, as they have been shown to bind and sequester the CKIs, p21 and p27, and by doing so they prevent p21 or p27 from binding and inhibiting cdk2-associated complexes (4–7). The association of p21 or p27 with cyclin D-cdk4 complexes creates a reservoir of CKIs, which can be mobilized in the presence of an antiproliferative response, to inhibit cdk2 activity and arrest cells in G1 phase (4). This association, however, serves another function, as p21 and p27 are required to assemble and stabilize cyclin D-cdk4, and under certain growth conditions, this association appears to be a non-inhibitory interaction (8, 9). In mouse embryonic fibroblasts (MEFs) derived from p21^−/−^-p27^−/−^ knockout mice, cyclin D-cdk4 complexes are not detected, while infection of these cells with retroviruses encoding either p21 or p27 restored cyclin D-cdk4 assembly (7). Other proteins also appear able to assemble cyclin D-cdk4 in the absence of p21 or p27, but this has not been well defined (10–12)

It is unclear whether both the catalytic and sequestration functions of cyclin D-cdk4 and cyclin D-cdk6 are required during normal proliferation or tumor progression, and the overexpression of cyclin D-cdk4 or cyclin D-cdk6 during tumor development would potentially increase both catalytic activity and the ability to sequester p27. While loss of both cdk4 and cdk6 leads to embryonic lethality, a remarkable amount of proliferation occurs before these animals die at E17.5, demonstrating that neither function is required during early fetal development or that cdk4 and cdk6 can be compensated by cdk1 or cdk2 (13–16). MEFs derived from these animals do not display any obvious defects in proliferation once in cycle. In contrast, serum-starved quiescent MEFs derived from either cdk4^−/−^ or cdk6^−/−^ embryos display a delay in re-entering the cell cycle upon serum stimulation, suggesting that some rate limiting function for cdk4 or cdk6 exists (13, 15). Loss of both cdk4 and cdk6 in cdk4^−/−^ - cdk6^−/−^ MEFs delays release from G0 even more (15). Whether this increased delay is due to the combined loss of cdk4/6 levels or to cdk4 and cdk6 specific functions is not known.

Cyclin D1^−/−^ mice are also viable (17), but display severe retinal and mammary gland defects. Due to the low levels of cyclin D2 and D3 in these tissues, loss of cyclin D1 causes the loss of both the catalytic and sequestration functions of cdk4 and cdk6 (17). These phenotypes can be rescued by the expression of the cyclin D1^K112E^ allele, which binds to cdk4 or cdk6 but cannot activate catalytic activity (18, 19). Cyclin D1-cdk4 complexes isolated from the retina of cyclin D1^K112E/K112E^ mice retain the ability to bind p27, suggesting that it is the kinase-independent functions that are involved in the development of this tissue (18, 19). However, MMTV-ErbB-2/cyclin D1^K112E/K112E^ mice do not develop breast cancer, suggesting that catalytic activity is specifically involved in tumor formation (18, 19) and is the downstream target of ErbB-2 oncogenic activation. It has been shown that lack of cyclin D-associated kinase activity reduces the progenitor population from which these ErbB-2 tumors will ultimately arise (20).

Cdk4 and cdk6 share approximately 70% amino acid sequence homology with each other, but only 40% homology with other identified cdks, placing them in a distinct subclass (21). As both cdk4 and cdk6 partner specifically with the D-type cyclins and phosphorylate the same residues in Rb, they have been considered functional homologues. This was in part confirmed in the knockout mice, as both single knockout cdk4^−/−^ and cdk6^−/−^ mice are viable, suggesting that functional redundancy existed in most tissues (13–16). However, some tissue-specific defects were observed. Cdk4-deficient mice developed type 1 diabetes-like phenotype, due to decreased numbers of pancreatic β cells, displayed reduced size compare to wild-type littermates, and had reduced fertility (13, 14). These defects were present even though cdk6 was expressed and bound to p27 in cdk4 null MEFs, arguing for a cdk4-specific role in certain tissues (13, 14). Cdk6-deficient mice, in contrast, had reduced thymus and spleen size due to reduced cellularity (15, 16). However, cdk4 was still present in cdk6 null T cells, was detected in complex with p27, and Rb was still phosphorylated (16), suggesting that in these tissues non-redundant functions existed as well.

In this study we sought to understand the contribution of the kinase-dependent and kinase-independent functions of cdk4 and cdk6 in epithelial cells, and to directly compare cdk4 and cdk6 in the same model system. While we found that overexpression of cdk4 caused cells to reenter cycle from G0 faster in a kinase-dependent fashion, overexpression of catalytically inactive cdk6 also accelerated G0 release. This cdk6 variant also caused the cell to overcome TGF-β-mediated growth arrest, proliferate in the absence of cyclin D-associated kinase activity and maintain cdk2 activity. Gel filtration analysis revealed the presence of a previously unidentified cyclin D-cdk6 dimer, which may contribute to the differential ability of cdk6 to affect p27 and cyclin D distribution in the cell and contribute to kinase-independent activity.

## EXPERIMENTAL PROCEDURES

### Plasmids

Constructs encoding human HA-cdk4^WT^, HA-cdk4^D158N^, HA-cdk6^WT^, HA-cdk6^D163N^, HA-cdk4^R24C^, HA-cdk4^D158N/R24C^, HA-cdk6^R31C^, cdk6^D163N/R31C^ and HA-cdk6^T177A^ and mouse cdk4^WT^, cdk4^K35M^, cdk4^R24C^, cdk4^K35M/R24C^ and cdk4^T172A^ were generated by direct cloning into pUHD 10-3 hygromycin (22) or using PCR mutagenesis.

### Cell Culture

Mv1Lu, Mv1Lu-tTA cells (4) and their derivatives were cultured in Minimal Essential Media supplemented with 10% fetal bovine serum. Media for the tetracycline-regulated cell lines also contained 2 μg/ml tetracycline, 500 μg/ml G418 and 300 μg/ml hygromycin. Tet-regulated lines expressing the cdk4 and cdk6 variants were generated by transfecting tTA cells with 25 μg of the appropriate DNA using Lipofectin (Invitrogen, Carlsbad, CA) according to the manufacturer’s protocol. Individual colonies generated during selection with hygromycin were grown in the absence of tetracycline and assayed for exogenous protein expression. To assay the effects of exogenous protein expression, the cell lines were grown in the presence of tetracycline (referred to in each figure as the *OFF* condition) or in the absence of tetracycline (referred to in each figure as the *ON* condition). Where indicated, cells were treated with 100 pM TGF-β (R&D Systems, Minneapolis, MN) for 24 hours. For cell cycle analysis, cells were harvested and analyzed by flow cytometry as previously described (8).

### Antibodies

The antibodies used in this study were as follows: anti-HA (Y-11), anti-cdk4 (C-22), anti-cyclin E (C-19), anti-cyclin D1 (M-20 and H-295), anti-cyclin D2 (C-17), anti-p27 (M-197, M-20 and C-19) and anti-cdk6 (C-21) (all from Santa Cruz, Santa Cruz, CA), anti-cyclin D1 (DCS-6) and anti-cyclin D2 (DCS-11) (both from Labvision, Fremont, CA), anti-p27 (C-57) and anti-pRb (G3-245) (both from BD Biosciences, San Jose, CA), anti-mouse p27, anti-mink cdk4, anti-mouse cdk4 and anti-mouse cdk2 (rabbit polyclonal, generous gifts from Dr. Joan Massagué), anti-cdk6 (DCS-90) anti-β actin (both from Sigma, St. Louis, MO), protein A-HRP (Millipore, Billerica, MA), goat anti-mouse-HRP (Bio-Rad, Hercules, CA), goat anti-mouse (Southern Biotech, Birmingham, AL) and goat anti-rabbit (Jackson Immunoresearch, West Grove, PA).

### Immunoblot Analysis, Immunoprecipitations and Kinase Assays

For immunoblot analysis, cells were lysed in TNT buffer (50 mM Tris, pH 8.0, 150 mM NaCl, 1 mM EDTA and 1% Triton X-100) supplemented with protease inhibitor cocktail V (Calbiochem, San Diego, CA) and 1 mM phenylmethylsulfonyl fluoride (PMSF). For all immunoprecipitations, cells were lysed in Tween-20 buffer as previously described (5). All imunoprecipitations, immunoblot analysis and *in vitro* kinase assays using recombinant Rb as a substrate were performed as previously described (8).

### Gel Filtration

Lysates were separated by size exclusion chromatography (gel filtration) as previously described (5, 23). Cells were lysed in Tween-20 buffer and the lysates were cleared by sonication followed by ultracentrifugation at 50,000 rpm. 4.5 mg of extract was fractionated on a Superdex 200 10/30 column (GE-Amersham, Piscataway, NJ) and fractions #19-34 were collected and analyzed from each run.

## RESULTS

### Exogenous cdk4 or cdk6 expression allows the cell to overcome TGF-β-mediated growth arrest

TGF-β treatment of mink lung epithelial (Mv1Lu) cells causes growth arrest in the G1 phase of the cell cycle (4). TGF-β induces expression of p15, which inhibits the catalytic activity of cyclin D-cdk4 and cyclin D-cdk6, and also displaces p27 from the cyclin D-cdk4 and cyclin D-cdk6 complexes, as p15 and p27 binding is mutually exclusive (22). Loss of the cdk4 and cdk6 sink permits the liberated p27 to bind and inhibit cdk2 complexes (22). Previously, it had been shown that overexpression of either cdk4 or cyclin D1 in Mv1Lu cells rendered them able to overcome TGF-β-mediated growth arrest (22–24). By increasing the cyclin D-associated-p27 sink, cdk4, cdk6, and cdk2 remained active and were able to evade inhibition by p15 and p27 (23).

Mv1Lu cells expressing different variants of HA-tagged human cdk4 (Tet-cdk4^WT^, Tet-cdk4^D158N^, Tet-cdk4^R24C^ and Tet-cdk4^D158N/R24C^), HA-tagged human cdk6 (Tet-cdk6^WT^, Tet-cdk6^D163N^, Tet-cdk6^R31C^ and Tet-cdk6^D163N/R31C^) or untagged mouse cdk4 (Tet-mcdk4^WT^, Tet-mcdk4^K35M^, Tet-mcdk4^R24C^ and Tet-mcdk4^K35M/R24C^) in a tetracycline-repressible manner were generated to assess the requirement for cdk4 and cdk6 kinase and kinase-independent activities during asynchronous proliferation. Cdk4^D158N^, cdk4^K35M^ and cdk6^D163N^ mutations specifically target the catalytic core, creating kinase inactive proteins that retain their ability to bind to cyclin D, p27 and p15 (25–27). The catalytically inactive proteins do not act as dominant-negatives in proliferating cells, as their overexpression does not alter cell cycle kinetics (26). In the presence of TGF-β, however, their expression might provide a larger sink for p27 or p15, potentially permitting the endogenous cdk4, cdk6 and cdk2 to remain active. Cdk4^R24C^ and cdk6^R31C^ mutants are catalytically active, can bind p27, but cannot be inhibited by p15 (27, 28). Therefore, in the presence of TGF-β, their expression could provide a sink for p27 to keep cdk2 inhibitor-free, but could also act to increase cdk4 or cdk6 catalytic activity. The double mutants, cdk4^D158N/R24C^, cdk4^K35M/R24C^ and cdk6^D163N/R31C^ lack catalytic activity, cannot be inhibited by INK4 proteins, but retain the ability to bind p27 (27). These are p27 sequestration-only mutants, and in these cells treated with TGF-β, we predicted that all cdk4 and cdk6 kinase activity would be inhibited by p15 expression.

To assess expression of the exogenous proteins, each line was grown in the presence of tetracycline (Tet) (hereafter indicated as the *OFF* condition in each figure, no exogenous protein expression) or in the absence of Tet (hereafter indicated as the *ON* condition in each figure, exogenous protein is expressed) for 24 h. and the resulting lysates were subjected to immunoblot analysis. As expected, removal of Tet induced exogenous gene expression (Fig. 1), when the cdk6 lines were probed with anti-cdk6 antibodies (Fig. 1A) and the human cdk4 lines were probed with either anti-HA or anti-cdk4 antibodies (Fig. 1B), or when the mouse cdk4 lines were probed with anti-mouse cdk4 antibodies, which do not recognize endogenous mink cdk4 (Fig. 1C). The cdk6 immunoblot was not probed with anti-HA antibodies because cdk6^D163N/R31C^ lacks an HA epitope tag. Also, expression of mouse cdk4^R24C^ resulted in several detectable degradation products of faster mobility (Fig. 1C). For each of the assays described herein, the data were reproduced with multiple independent clones of each line, but the results from only one clone are shown for simplicity.

**Figure 1.**
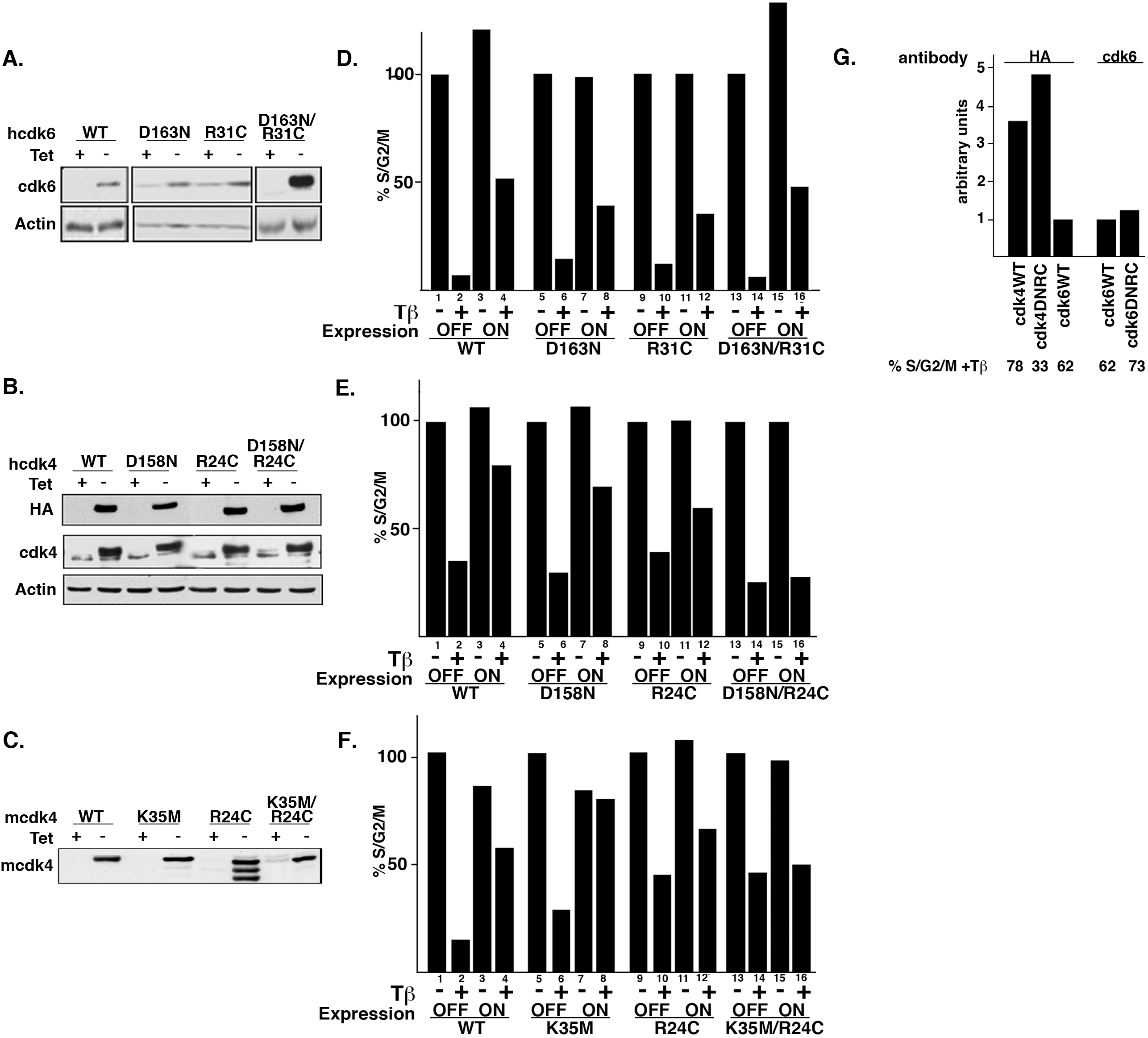
Expression of sequestration-only cdk6, but not cdk4, allows cells to overcome TGF-β-mediated growth arrest. Each cell line was grown for 20 hours in the presence (+) or absence (-) of tetracycline (Tet) and either harvested for immunoblot analysis for detection of the exogenous protein, treated with TGF-β for an additional 24 hours and analyzed for DNA content by flow cytometry. **A**. Lysates from Tet-hcdk6^WT^, Tet-hcdk6^D163N^, Tet-hcdk6^R31C^ and Tet-hcdk6^D163N/R31C^ cells grown +/- Tet were analyzed for cdk6 expression by direct immunoblot using anti-cdk6 antibodies. **B.** Tet-hcdk4^WT^, Tet-hcdk4^D158N^, Tet-hcdk4^R24C^ and Tet-hcdk4^D158N/R24C^ cells were treated and analyzed similar to A. Lysates from the cells were analyzed by immunoblot using anti-cdk4 and anti-HA antibodies. **C.** Tet-mcdk4^WT^, Tet-mcdk4^K35M^, Tet-mcdk4^R24C^ and Tet-mcdk4^K35M/R24C^ were treated and analyzed similar to A. The immunoblot was probed with anti-mouse cdk4 antibodies, which does not cross-react with the endogenous mink cdk4. The results shown are representative of at least three independent experiments. **D**. Tet-cdk6 cells grown OFF or ON and treated +/- TGF-β were analyzed for DNA content by flow cytometry. The percentage of cells in S/G2/M in each line treated + Tet/- TGF-β was set to 100%. **E.** Tet-cdk4 cells grown OFF or ON and treated +/- TGF-β were analyzed for DNA content by flow cytometry similar to D. **F.** Tet-mcdk4 cells grown OFF or ON and treated +/- TGF-β were analyzed for DNA content by flow cytometry similar to D. **G.** The relative expression levels of cdk4^WT^, cdk4^D158N/R24C^ and cdk6^WT^ were determined by densitometry analysis of an immunoblot probed with anti-HA antibodies. The relative expression levels of cdk6^WT^ and cdk6^D163N/R31C^ were determined by densitometry analysis of an immunoblot probed with anti-cdk6 antibodies. The percentage of S/G2/M cells from the same samples treated with TGF-β is also shown.

To determine the effect of exogenous cdk6 expression on cell cycle progression, the cell lines were grown in the OFF or ON conditions for 24 h. and TGF-β was added for an additional 24 h. before the cells were harvested for cell cycle analysis. The percentage of Tet-cdk6^WT^ cycling cells (S/G2/M content) seen in the OFF/-TGF-β condition was set to 100% (Fig. 1D, lane 1). TGF-β arrested these cells, resulting in a reduced S/G2/M content (Fig. 1D, lane 2). A similar pattern of TGF-β-mediated arrest was seen with all cells grown in the OFF condition (Fig. 1D, lanes, 5, 6, 9, 10, 13, 14). Exogenous protein expression did not affect proliferation as cells grown in the ON condition (Fig. 1D, lanes 3, 7, 11, 15) had similar S/G2/M content as cells grown in the OFF condition (Fig. 1D, lanes, 1, 5, 9, 13). When the Tet-cdk6^WT^, cdk6^D163N^ and cdk6^R31C^ cells were grown in the ON condition and treated with TGF-β, however, we found that expression of exogenous cdk6 led to a partial recovery from TGF-β-mediated growth arrest (Fig. 1D, lanes 4, 8, 12). When the Tet-cdk6^D163N/R31C^ cells were grown in ON condition and treated with TGF-β, they were also able to partially overcome TGF-β-mediated growth arrest (Fig. 1D, lane 16). As cdk6^D163N/R31C^ cannot bind p15 and does not have catalytic activity (27), its only functions should be kinase-independent and might involve sequestering p27. In the presence of TGF-β, p15 should bind and inhibit the endogenous cdk4 and cdk6, suggesting that these cells might be proliferating in the absence of cdk4 and cdk6 kinase activity. We extended this assay to 48 h. of TGF-β treatment, cells numbers were enumerated by counting with a hemocytometer and we found that both cdk6^WT^ and cdk6^D163N/R31C^ were still competent to rescue proliferation in the presence of TGF-β over multiple rounds of division (data not shown).

We next assayed the ability of the human cdk4 cell lines to overcome TGF-β-mediated growth arrest. All of the cdk4 lines were arrested in G1 and had reduced S/G2/M content when grown in the OFF condition and treated with TGF-β (Fig. 1E, lanes 2, 6, 10, 14). Expression of the exogenous cdk4 variants in the ON condition did not affect cell cycling and similar S/G2/M content was detected (Fig. 1E, lanes 3, 7, 11, 15), and like their cdk6 counterparts, expression of cdk4^WT^, cdk4^D158N^ or cdk4^R24C^ allowed the cells to partially overcome TGF-β-mediated growth arrest when grown in the ON/+TGF-β condition (Fig. 1E, lanes 4, 8, 12). Unexpectedly, expression of the sequestration-only cdk4^D158N/R24C^ mutant did not rescue TGF-β-mediated growth arrest, as a decrease in S/M/G2 content was detected (Fig. 1E, lane 16).

To verify that this inability to overcome TGF-β-mediated growth arrest was not due to a perturbation of the cdk4 structure by the introduction of the D158N/R24C mutation, we analyzed the ability of a similar panel of mouse cdk4 mutant cells to overcome TGF-β-mediated growth arrest (Fig. 1F). Similar to the results from the human cdk4 cell lines, the sequestration-only mouse cdk4, cdk4^K35M/R24C^, did not support proliferation in the presence of TGF-β (Fig. 1F, lane 16), suggesting that cdk4 and cdk6 differed in their abilities to overcome TGF-β-mediated growth arrest. To rule out that the sequestration-only cdk6 mutant was expressed to a higher level than the sequestration-only cdk4 mutant, we compared the level of exogenous protein expression between the human cdk4 and cdk6 lines and found that cdk4 proteins were expressed at levels greater than the cdk6 proteins (Fig. 1G), The level of expression did not correlate with the ability to overcome TGF-β-mediated growth arrest, as seen by the percentage of S/G2/M cells harvested from the same sample. This suggests that the inability of cdk4^D158N/R24C^ to rescue proliferation was not due to a reduced level of expression.

### Expression of cdk6^D163N/R31C^ rescues cdk2 activity in the presence of TGF-β

In the presence of TGF-β, endogenous cdk4 and cdk6 kinase activity should be inhibited by p15 expression. As the cdk6^D163N/R31C^ mutant is catalytically inactive, we hypothesized that cdk6^D163N/R31C^ cells might be proliferating in the absence of cdk4 and cdk6 kinase activity. We also thought that expression of cdk6^D163N/R31C^ might provide a sink to sequester liberated p27 and cdk2 kinase activity might be restored in the presence of TGF-β treatment. To test this hypothesis, we examined p27-associated and cdk2-associated kinase activity using Rb *in vitro* kinase assays. Cdk4- and cdk6-expressing cells from the four conditions described above (OFF, OFF/+TGF-β, ON, ON/+TGF-β) were harvested to generate lysates. p27- and cdk2-associated complexes were isolated by immunoprecipitation with anti-p27 or anti-cdk2 antibodies, respectively, and mixed with recombinant Rb protein and γ-P^[32]^ATP. While p27 immunoprecipitates contain cdk4, cdk6 and cdk2, p27-associated kinase activity has been shown to be due exclusively to cdk4 and cdk6 (5, 8) and is a marker of cyclin D-associated kinase activity. In Tet-cdk6^WT^ lysates from cells grown in the OFF condition, both p27- and cdk2-associated kinase activities were detected (Fig. 2A, lane 1). In Tet-cdk6^WT^ lysates generated from cells in the OFF/+TGF-β condition, both p27-associated and cdk2-associated kinase activities were repressed (Fig.2A, lane 2), consistent with TGF-β’s inhibition of cdk4, cdk6 and cdk2 kinase activity (4). In lysates from Tet-cdk6^WT^ cells grown in the ON condition, p27- and cdk2-associated kinase activity was detected (Fig. 2A, lane 3), and expression of exogenous cdk6 did not significantly increase either activity. In lysates from Tet-cdk6^WT^ cells grown in the ON/+ TGF-β condition, p27- and cdk2-associated kinase activities were retained (Fig. 2A, lane 4), suggesting that the increase in the cdk6 expression permitted cdk4, cdk6 and cdk2 to remain active in the presence of TGF-β.

**Figure 2.**
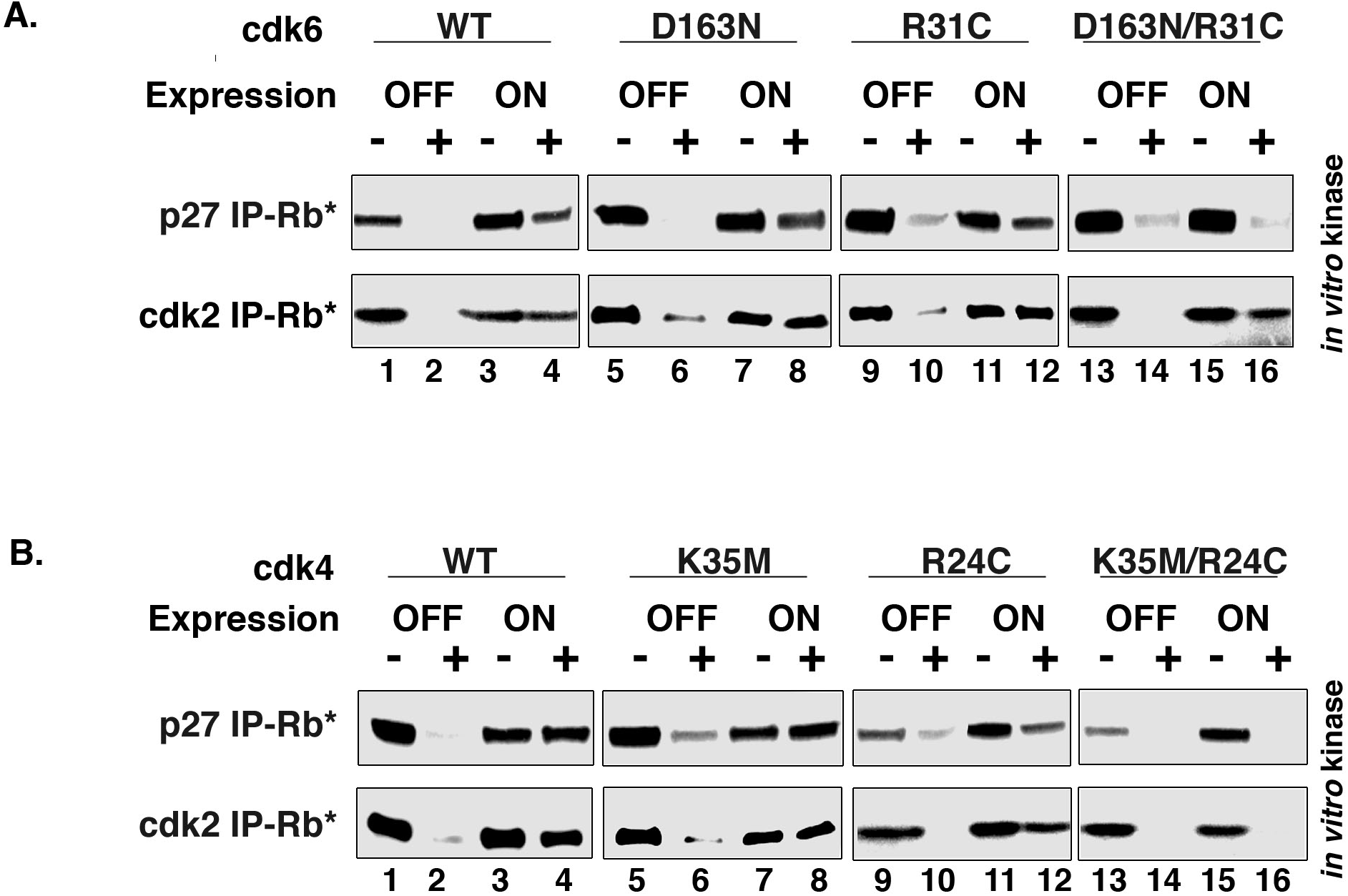
Cdk6^D163N/R31C^ expression supports cdk2 catalytic activity in the presence of TGF-β. Lysates from cdk6 cells **A**. or mcdk4 cells **B.** grown in the presence (OFF) or absence (ON) of Tet and treated +/- TGF-β were subjected to immunoprecipitation with either anti-p27 or anti-cdk2 antibodies and used in Rb *in vitro* kinase assays. Note, Rb*=phosphorylated Rb. The results shown are representative of three independent experiments.

Similar results were seen with Tet-cdk6^D163N^ (Fig. 2A, lanes 5-8) and cdk6^R31C^ (Fig. 2A, lanes 9-12) lysates, where p27-associated and cdk2-associated kinase activities were retained in the ON/+TGF-β condition, consistent with the proliferation of these cells (Fig. 1D, lanes 8, 12).

When lysates from Tet-cdk6^D163N/R31C^ cells, grown in the OFF condition, were assayed for kinase activity, p27- and cdk2-associated kinase activities were detected (Fig. 2A, lane 13), which were then inhibited by the addition of TGF-β (OFF/+TGF-β) (Fig. 2A, lane 14). Lysates from cells grown in the ON condition also had p27- and cdk2-associated activity (Fig. 2A, lane 15), demonstrating that the expression of the catalytically inactive cdk6 variant did not affect total cyclin D-associated kinase activity. However, lysates from cells grown in the ON/+ TGF-β lost p27-associated kinase activity but retained cdk2-associated kinase activity (Fig 2A, lanes 15 and 16). This suggested that TGF-β treatment was sufficient to inhibit all cdk4 and cdk6 activity, but the expression of the cdk6^D163N/R31C^ was capable of maintaining cdk2 kinase activity.

We performed similar experiments with lysates generated from cells expressing the cdk4 mutants. Lysates from cells expressing cdk4^Wt^(Fig. 2B, lanes 1-4), cdk4^K35M^ (Fig. 2B, lanes 5-8) and cdk4^R24C^ (Fig. 2B, lanes 9-12), all exhibited similar patterns of activity: p27 and cdk2-associated activity was detected in the OFF and ON conditions (Fig. 2B, lanes 1, 3, 5, 7, 9, 11). This activity was inhibited by TGF-β treatment in the OFF/+TGF-β condition (Fig. 2B, lanes 2, 6, 10). The expression of cdk4^WT^, cdk4^K35M^, or cdk4^R24C^ variants in the ON/+TGF-β condition caused p27- and cdk2-associated activity to be maintained (Fig. 2B, lanes 4, 8, 12), consistent with the ability to overcome TGF-β-mediated growth arrest seen with these cells (Fig. 1D).

When lysates from cdk4^K35M/R24C^ cells grown in the OFF condition were analyzed, p27- and cdk2-associated kinase activities were detected, which were inhibited by the addition of TGF-β (OFF/+TGF-β) (Fig. 2B, lanes 13, 14). When these cells were grown in the ON condition (Fig. 2B, lane 15), p27-associated kinase activity was still detected, suggesting that this cdk4 variant did not act as a dominant negative mutant. However, lysates from cdk4^K35M/R24C^ cells grown in the ON/+ TGF-β condition, did not have either p27- or cdk2-associated kinase activity (Fig. 2B, lane 16). Similar to expression of the cdk6^D163N/R31C^ variant, expression of cdk4^K35M/R24C^ in the presence of TGF-β resulted in the loss of all cdk4- and cdk6-associated kinase activity, but expression of the cdk4^K35M/R24C^ mutant was unable to restore cdk2-associated kinase activity (Fig. 2B, lane 16). This data suggested that the cdk6^D163N/R31C^ cells were able to proliferate in the absence of cdk4- and cdk6-associated kinase activity because cdk2 remained active in the presence of TGF-β (Fig. 2A, lane 16), while cdk4^K35M/R24C^ expressing cells did not maintiain cdk2 kinase activity and thus were unable to proliferate in the presence of TGF-β.

### Cdk4 and cdk6 accelerate exit from G0 by two different mechanisms

Previous studies had suggested that the catalytic function of cdk4 or cdk6 might be specifically required during the G0 to G1 transition, and loss of cdk4 or cdk6 in null MEFs causes a retardation in the exit from quiescence (13, 15). To determine whether overexpression of cdk4 or cdk6 had any affect on the G0-G1 transition, Tet-cdk4^WT^, Tet-cdk4^D158N/R24C^, Tet-cdk6^WT^ and Tet-cdk6^D163N/R31C^ cells were grown to visible contact arrest in the OFF or ON condition and maintained in a quiescent state, with serum replenished every other day. After 5 days, the cells were replated at a lower density, harvested at different time points post-release and subjected to DNA content analysis by flow cytometry in order to follow cell cycle reentry.

First, we analyzed Tet-cdk4^WT^ cells arrested and replated in the OFF condition (Fig. 3A). These cells exit the G0 state slowly and by 24 h. post-release 30% of the cells had exited G1 and an increase in the S/G2/M content could be detected (Fig. 3A). In contrast, 30% of Tet-cdk4^WT^ cells arrested and replated in the ON condition had increased their S/G2/M content by 16 h. post-release. The S/G2/M content peaked at 19 h. post-release and by 24 h., it was only 25%, indicating that the cells had completed one cell cycle and were re-entering G1 phase (Fig. 3A).

**Figure 3.**
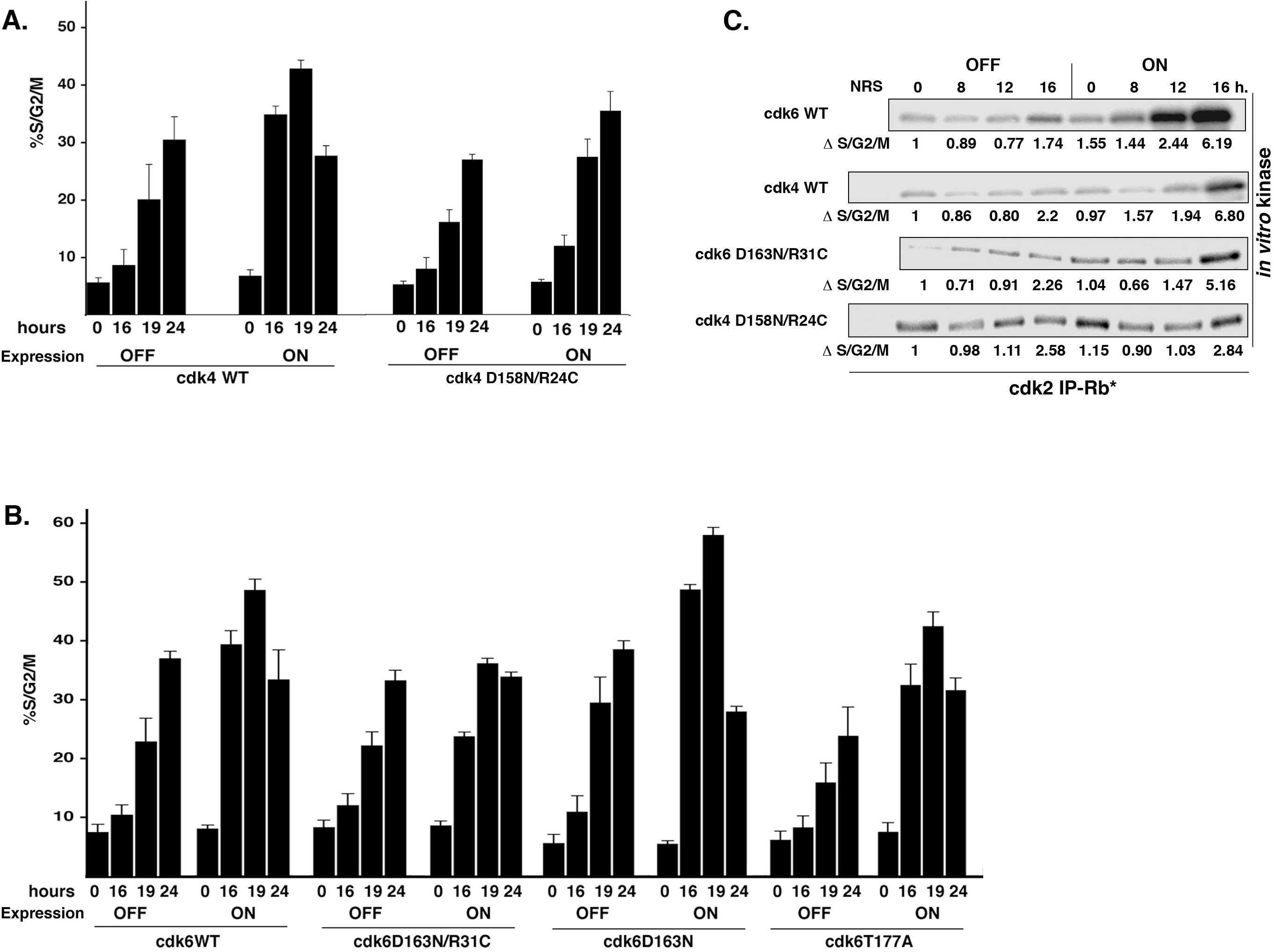
Cdk4 catalytic activity and cdk6 non-catalytic activity accelerates the exit from G0 and correlates with cdk2 activity. **A.** Tet-cdk4^WT^ and Tet-cdk4^D158N/R24C^ cells were grown to confluence in the presence (OFF) or absence (ON) of Tet, maintained in a quiescent state in the presence of serum for 5 days and then stimulated to proliferate by replating at a lower density. Cells were harvested at 0, 16, 19 and 24 hours post-release and analyzed for S/G2/M content by flow cytometry. **B**. Tet-cdk6^WT^, Tet-cdk6^D163N/R31C^, Tet-cdk6^D163N^ and Tet-cdk6^T177A^ cells were treated and analyzed as in A. The results shown are the average of at least three independent experiments with standard deviation. **C.** Tet-cdk4^WT^, Tet-cdk6^WT^, Tet-cdk4^D158N/R24C^ and Tet-cdk6^D163N/R31C^ cells were grown to confluence and replated in the presence (OFF) or absence (ON) of Tet.. Lysates from cells harvested at 0, 8, 12 and 16 hours post-release were immunoprecipitated with anti-cdk2 antibodies and used in Rb in vitro kinase assays. Fixed cells from the same samples were analyzed for DNA content by flow cytometry. The percentage of S/G2/M cells was adjusted so that the value at time 0 in the presence of Tet was equal to 1. The S/G2/M population of the other samples from each line was then expressed as a change in population relative to time 0, denoted as Δ S/G2/M.

Tet-cdk4^D158N/R24C^ cells arrested in the OFF condition exited G0 with kinetics similar to those seen with the cdk4^WT^ cells. An increase in the S/G2/M content was seen by 19 h. post-release that continued to increase by 24 h. post release (Fig. 3A). When these cells were maintained and replated in the ON condition, the presence of the exogenous cdk4^D158N/R24C^ did not cause cells to exit the G0 phase faster and the rate of increase of the S/G2/M population mirrored that seen with cells grown in the OFF condition (Fig. 3A). Overexpression of two other catalytically inactive cdk4 mutants (mcdk4^T172A^ (29) and mcdk4^K35M^, data not shown) also failed to accelerate G0 release, suggesting that exogenous cdk4 fulfills a rate-limiting kinase-dependent function.

We tested the ability of cdk6 to affect the exit from G0. Tet-cdk6^WT^ cells grown in the OFF condition steadily exited G0, while cells grown in the ON condition exited G0 more rapidly, and a dramatic increase in the S/G2/M content was seen as early as 16 h. post-release (Fig. 3B). The increase in the S/G2/M content peaked by 19 h. and decreased by 24 h., suggesting that these cells has completed one cell cycle and were re-entering G1 phase (Fig. 3B). Tet-cdk6^D163N/R31C^ cells grown in the OFF condition exited G0 with kinetics similar to the cdk6^WT^ cells (Fig. 3B). Surprisingly, Tet-cdk6^D163N/R31C^ cells grown in the ON condition exited G0 more rapidly when compared to the cells grown in the OFF condition, with 30% of the cells exhibiting a significant increase in their S/G2/M content by 16 h. post-release (Fig. 3B). This peaked at 19 h post-release and began to decrease by 24 h (Fig. 3B), suggesting that cdk6 could accelerate G0 exit in a kinase-independent manner.

We tested two other catalytically inactive mutants, cdk6^D163N^ and cdk6^T177A^ in our release from contact assay. The Tet-cdk6^T177A^ cells express a cdk6 mutant that cannot be phosphorylated by CAK and is similar to cdk4^T172A^ (30). When these lines were arrested and replated in the OFF condition, the S/G2/M content slowly increased over 24 h., consistent with the slow exit from G0 seen with the Tet-cdk6^WT^ and Tet-cdk6^D163N/R31C^ cells grown in the OFF condition (Fig. 3B). In contrast, when both of these lines were arrested and replated in the ON condition, there was a dramatic increase in their S/G2/M content by 16 h post-release (Fig. 3B), suggesting that the expression of catalytically inactive cdk6 was sufficient to accelerate exit from G0 (Fig. 3B).

We also adjusted the percentage of S/G2/M cells from arrested and restimulated conditions so that the value at time 0 in the OFF condition was equal to 1 (Fig. 3C). The S/G2/M population of the other samples from each line was then expressed as a change in population relative to time 0, denoted as Δ S/G2/M. For all of the cells grown in the OFF condition, the Δ S/G2/M by 16 h. was approximately 2 fold (Fig. 3C). By 16 h., the Δ S/G2/M for the cdk6^WT^, cdk4^WT^, or cdk6^D163NR31C^ grown in the ON conditions increased to a similar extent, between 5-7 fold, suggesting that the rate-limiting advantage seen by overexpression of catalytically active cdk4 and cdk6 or catalytically inactive cdk6 was roughly equivalent (Fig. 3C). The Δ S/G2/M for cdk4^D158N/R24C^ was that same in the ON and OFF condition, consistent with the lack of acceleration seen in the presence of this catalytically inactive mutant (Fig. 3C).

Cdk2 kinase activity becomes detectable at the G1-S phase border, and lysates from the arrested and restimulated cells were analyzed for cdk2-associated Rb kinase activity *in vitro* as a marker of G0-release. Cdk2 kinase was inactive in all G0 arrested cell lines (Fig. 3C, 0h.) as seen by Rb *in vitro* kinase assay. Between 0 h. and 16 h. post-release, cdk2-associated kinase activity was still not detectable in the Tet-cdk6^WT^, Tet-cdk4^WT^, or Tet-cdk6^D163N/R31C^ cells arrested and replated in the OFF condition. However, cdk2 activity did increase dramatically in cells arrested and replated in the ON condition (Fig. 3C). This was in contrast to what was seen with the Tet-cdk4^D158N/R24C^ cells arrested and replated in the OFF or ON conditions between 0 h. and 16 h (Fig. 3C), where cdk2-associated kinase activity remained constant and inactive in both conditions (Fig. 3C).

### Both cdk4 and cdk6 function as a p27 sink

Data suggested that expression of cdk6^D163N/R31C^ was sufficient to both render cells resistant to TGF-b-mediated growth arrest and accelerate release from G0 phase, while the homologus mutation in cdk4, cdk4^D158N/R24C^, was not. Cdk6^D163N/R31C^ did not appear to be expressed to a higher level than the cdk4 variant (Fig. 1G), and by indirect immunofluorescence we found that _both cdk4_D158N/R24C _and cdk6_D163N/R31C _were_ expressed in similar subcellular patterns, in both the nuclear and the cytoplasm (data not shown), suggesting that this also could not account for their differential abilities. We hypothesized that cdk6 might sequester p27 more efficiently than cdk4 could. We examined the composition of complexes isolated from Tet-cdk4^WT^ and Tet-cdk6^WT^ cells grown in the ON condition and incubated with or without TGF-β using co-immunoprecipitation-immunoblot analysis with anti-cyclin D1, anti-cyclin D2, anti-p27 and anti-HA antibodies (Fig. 4A). Cdk4 and cdk6 were present at comparable levels in cyclin D1, cyclin D2 and p27 immunoprecipitates from cdk4^WT^ and cdk6^WT^ cells grown in the absence or presence of TGF-β, as seen with anti-HA antibodies (Fig. 4A, panels 1-3), suggesting that both exogenous cdk4 and cdk6 were able to associate efficiently with cyclins D1 and D2 and p27. Immunoprecipitation with anti-HA antibodies, followed by HA immunoblot analysis, demonstrated that HA-tagged cdk4 and cdk6-associated complexes were recovered to a similar extent in this assay (Fig. 4A, panel 4). Cyclin D1, cyclin D2 and p27 levels were relatively unchanged in all of the conditions surveyed (Fig. 4A, panels 6, 7 and 8, Fig. 4B, panels 10 and 11), and a direct immunoblot with HA antibodies confirmed that endogenous cdk4 levels were in fact greater than those seen with cdk6 (Fig. 4A, panel 9). From this data, we concluded that there was no difference between exogenous cdk4^WT^ and cdk6^WT^ in the ability to bind to cyclin D1, cyclin D2 and p27.

**Figure 4.**
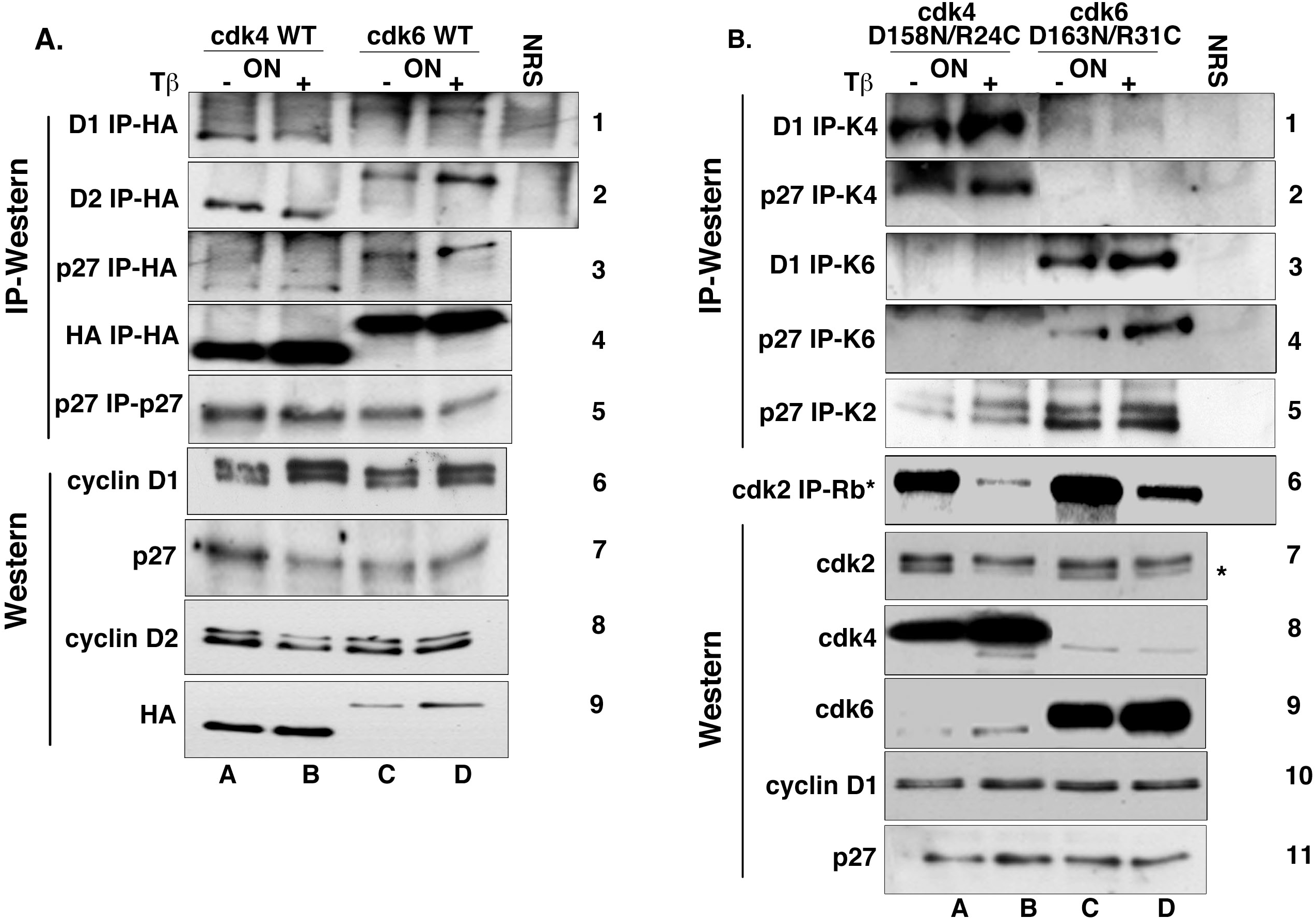
Cdk4 and cdk6 are similar p27 sinks. Each cell line was grown in the absence of Tet (ON) and treated with or without TGF-β for 24 h. **A.** Lysates from TGF-β treated Tet-cdk4^WT^ (lanes A, B) and Tet-cdk6^WT^ (lanes C, D) cells were immunoprecipitated with anti-cyclin D1 (panel 1), anti-cyclin D2 (panel 2), anti-p27 (panels 3, 5) and anti-HA (panel 4) antibodies and immunoblotted with anti-HA or anti-p27 antibodies. Immunoblots of whole cell lysates were analyzed for cyclin D1 (panel 6), p27 (panel 7), cyclin D2 (panel 8) and HA-cdk4/6 (panel 9) expression. **B.** Lysates from TGF-β treated Tet-cdk4^D158N/R24C^ (lanes A, B) and Tet-cdk6^D163N/R31C^ (lanes C, D) cells were immunoprecipitated with anti-cyclin D1 (panels 1,3) and anti-p27 (panels 2, 4 and 5) antibodies and immunoblotted with anti-cdk4, anti-cdk6 or anti-cdk2 antibodies. A sample from the same lysates was immunoprecipitated with anti-cdk2 antibodies and used in Rb *in vitro* kinase assays (panel 6). Note, Rb*=phosphorylated Rb. Immunoblots of whole cell lysates were probed withanti-cdk2 (panel 7), anti-cdk4 (panel 8), anti-cdk6 (panel 9), anti-cyclin D1 (panel 10) or anti-p27 (panel 11) antibodies. Note in panel 9, *= phosphorylated cdk2. The results shown are representative of three independent experiments.

We wanted to examine the composition of total cdk4- or cdk6-associated complexes, which we assumed would be a mixture of endogenous and exogenous protein, to determine what effect exogenous expression would have on complex formation *in vivo*. Increased expression of cdk4^D158N/R24C^ and cdk6^D163N/R31C^ was verified by direct immunoblot analysis of lysates with anti-cdk4 and anti-cdk6 antibodies, respectively (Fig. 4B, panels 8 and 9). Complexes isolated from cdk4^D158N/R24C^ and cdk6^D163N/R31C^ cells grown in the ON condition and treated with or without TGF-β were analyzed by co-immunoprecipitation-immunoblot analysis with anti-cyclin D1 and anti-p27 antibodies (Fig. 4B). Cdk4 was detected in both cyclin D1 and p27 immunoprecipitates from cdk4^D158N/R24C^ cells grown in the absence and presence of TGF-β (Fig 4B, lanes A and B, panels 1 and 2), demonstrating that this complex was able to resist the expected destruction by TGF-β-induced p15. Cdk6 was detected in both cyclin D1 and p27 immunoprecipitates from cdk6^D163N/R31C^ cells grown in both conditions (Fig.3B, lanes C and D, panels 3 and 4). An increase in the amount of p27-cdk6 complex formation was seen in the presence of TGF-β, suggesting that liberated p27 might be associating with and increasing the assembly of cdk6 complexes. We observed that the exogenous kinase expressed in each cell type became the dominant kinase, with very little cyclin D-cdk6-p27 complexes being detected in the cdk4^D158N/R24C^ cells and vice versa (Fig. 4B, panels 1-4, lanes A-D). When p27 immunoprecipitates from cdk4^D158N/R24C^ lysates were probed with cdk2 antibodies, the expected TGF-β-dependent increase in asscociated cdk2 was seen (Fig.), suggesting that liberated p27 also bound to cdk2. The expression of cdk4^D158N/R24C^ in the presence of TGF-β did not support cdk2-associated kinase activity as measured by Rb *in vitro* kinase assays from cdk2 immunoprecipitates (Rb*) and by loss of the faster migrating, phosphorylated form (cdk2*) by immunoblot analysis (Fig. 4B, panels 6 and 7). The increased association of p27 with cdk2 presumably resulted in cdk2 inactivity. However, the amount of p27-cdk2 seen in the cdk6^D163N/R31C^ lysates was unchanged, and cdk2-associated kinase activity was maintained (Fig. 4B, panels 6 and 7). Our data suggest that while both cdk6^D163N/R31C^ and cdk4^D158/NR24C^ were competent to associate with cyclin D and p27 (Fig. 4B), only overexpression of cdk6 prevented the TGF-β-mediated increase in p27’s association with cdk2.

### Cdk6 complex formation determined by gel filtration chromatography

Gel filtration chromatography allowed us to further analyze the distribution of cdk4, cdk6 and associated proteins in lysates from continuously proliferating (A), contact arrested (G0) and TGF-β treated Tet-cdk6^WT^ cells grown in the OFF or ON condition. In A cells grown in the OFF condition, cdk4 was present equally in three distinct pools, which have been described previously: 400 kd, 200 kd and 66 kd (Fig. 5A, fractions 20-22, 25-27 and 30-32, respectively) (23,31–34). The 400 kd pool contains cdk4 complexed with molecular chaperones such as cdc37 and Hsp 90 (23,33,34). The 200 kd pool contains cyclin D-cdk4-p27 and cyclin D-cdk6-p27 complexes. In continuously proliferating cells, this is the only complex that has cdk4-, cdk6-, cyclin D- or p27-associated kinase activity (8,31,32). The small 66 kd pool contains cdk4 and cdk6 bound by Ink4 inhibitor proteins under certain conditions, monomeric cdk4, monomeric cdk6 and monomeric cyclin D (8,32,33). In the Tet-cdk6^WT^ cells grown in the OFF condition (Fig. 5A), the majority of cdk6 was found in the 66 kd pool (Fig. 5A, panel 2, fractions 30-32) and all of the p27 was found in the 200 kd fraction (Fig. 5A, panel 3, fractions 25-27). Cdk4 was detected in all three pools (Fig. 5A, panel 1, fractions 20-22, 25-27 and 30-32).

**Figure 5.**
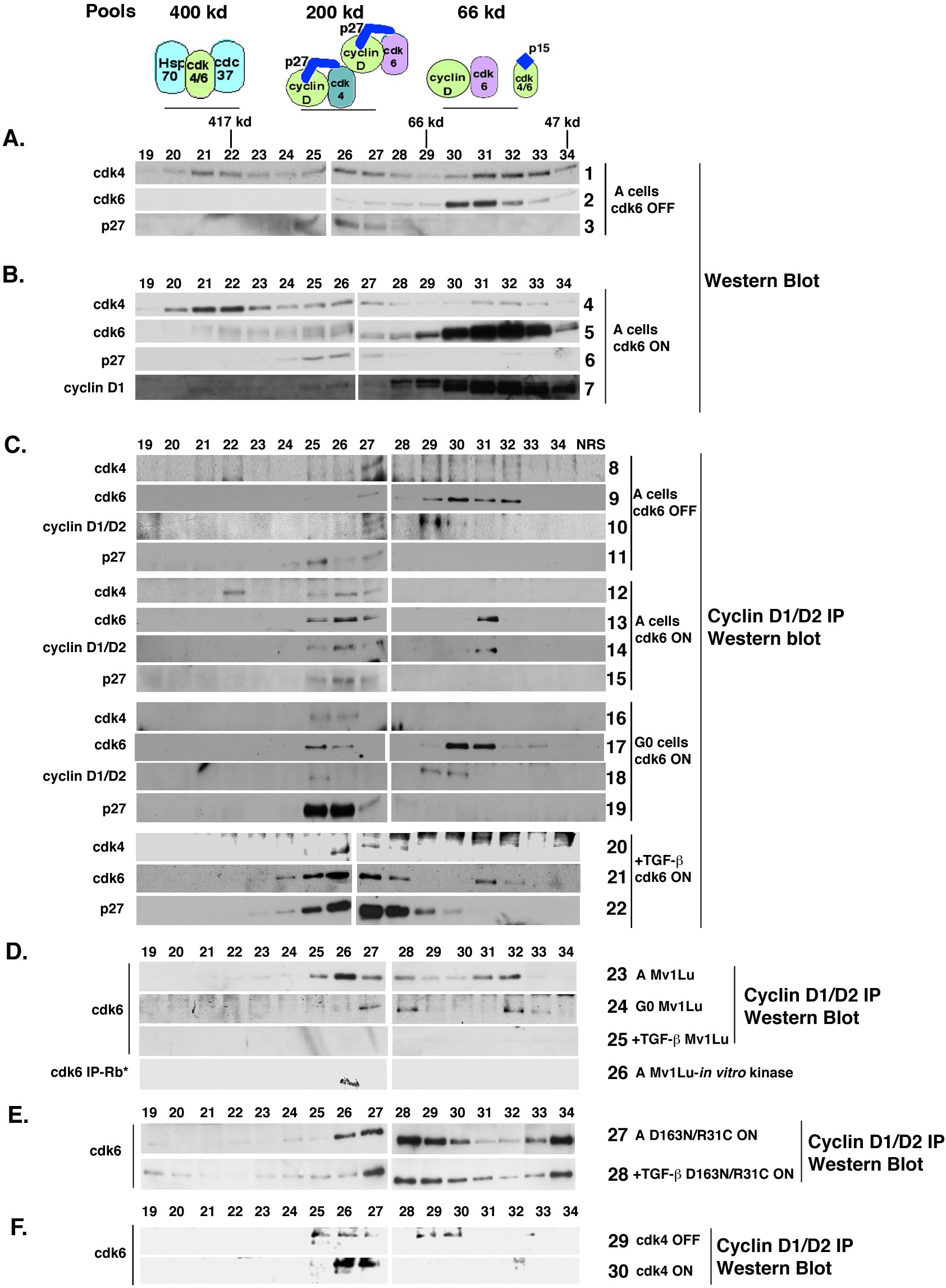
Cdk6 is able to form dimers with cyclin D1/D2 and increased cdk6 expression leads to increased formation of cyclin D-cdk6-p27 complexes. Tet-cdk6^WT^ cells were grown in the presence (OFF) or absence (ON) of Tet and, where indicated, either grown to confluence (G0) or treated with TGF-β. TGF-β was added to continuously proliferating cells. Lysates from these cells were separated by gel filtration on a Superdex 200 column and the resulting fractions were subjected to direct immunoblot analysis or immunoprecipitation-immunoblot analysis. **A.** Fractions from proliferating (A) Tet-cdk6WT cells grown in the presence of Tet (cdk6 OFF) were subjected to direct immunoblot analysis using anti-cdk4, anti-cdk6 and anti-p27 antibodies. **B.** Fractions from A Tet-cdk6WT cells grown in the absence of Tet (cdk6 ON) were analyzed by direct immunoblot with anti-cdk4, anti-cdk6, anti-p27 and anti-cyclin D1 antibodies. **C.** Fractions from A, G0 and TGF-β treated Tet-cdk6WT cells grown in the presence or absence of Tet were simultaneously immunoprecipitated with anti-cyclin D1 and anti-cyclin D2 antibodies. The resulting complexes were analyzed by immunoblotting with anti-cdk6, anti-cdk4, anti-cyclin D1, anti-cyclin D2 and anti-p27 antibodies. **D.** Lysates from G0 and TGF-β treated Mv1Lu cells were seperated by gel filtration and the resulting fractions were processed as described in C. An immunoblot was probed with anti-cdk6 antibodies. Fractions from A Mv1Lu cells were also immunoprecipitated with anti-cdk6 antibodies and used in an Rb *in vitro* kinase assay. **E.** Fractions from A and TGF-β treated Tet-cdk6D163N/R31C cells grown in the absence of Tet (ON) were subjected to cyclin D1/D2 immunoprecipitation and used in an anti-cdk6 immunoblot. **F.** Tet-mcdk4WT cells grown in the presence or absence of Tet for 24 h. were separated by gel filtration and the resulting fractions were processed as in C. An immunoblot from the resulting complexes was analyzed with anti-cdk6 antibodies. The location of the molecular weight standards is indicated on the top of part A. The results shown are representative of two independent experiments.

We then compared this profile to continuously proliferating Tet-cdk6^WT^ cells grown in the ON condition (Fig. 5B, panels 4-7). Increased cdk6 expression in A cells increased the percentage of cdk4 present in the 400 kd pool, but did not block formation of the 200 kd or 66 kd pools (Fig. 5B, panel 4, fractions 20-22, 25-27 and 30-32). Overexpression of cdk6 increased the formation of a 200kd cdk6 complex (Fig. 5B, panel 5, fractions 25-27). This was in contrast to cells grown in the OFF condition (A cells, cdk6 OFF), where there was little detectable cdk6 in the 200 kd fractions (Fig. 5A, panel 2, fractions 25-27). Increased cdk6 expression did not affect p27 distribution, which was still present in the 200 kd fractions (Fig. 5A and B, panels 3 and 6, fractions 25-27). A majority of cyclin D1 was located in the 66 kd pool, with a small amount present in the 200 kd pool, consistent with previous findings (Fig. 5B, panel 7, fractions 30-32 and 25-27, respectively) (8). Thus, our data suggested that increased expression of exogenous cdk6 did increase the amount of cdk6-associated with cyclin D and p27 in the 200 kd pool, although the majority of cdk6 did remain in the 66 kd pool.

To determine which gel filtration fractions contained cdk6 bound to the D-type cyclins, we co-immunoprecipitated cyclin D1/cyclin D2 complexes from the Tet-cdk6^WT^ cells. Mv1Lu cells do not express cyclin D3 (22). Previous studies had demonstrated that in Mv1Lu and primary T cells, cyclin D1-cdk4 complexes are found only in the 200 kd pool and contain p27, and we observed a similar result (Fig. 5C, panels 8 and 11, fractions 25-27) (8,23,31). This was consistent with observations that cyclin D cannot complex with cdk4 in the absence of p27 or another assembly factor. In A cells grown in the OFF condition, we found that cdk6 was bound to cyclins D1 and D2 in the 200 kd pool (Fig. 5C, panel 9, fractions 25-27), indicating the existence of cyclin D-cdk6-p27 ternary complexes. Unexpectedly, we detected cdk6 in the range of the 66 kd complex as well (Fig. 5C, panel 9, fractions 30-32), suggesting that cdk6 was able to dimerize with cyclin D in the absence of p27. p27-cyclin D1/D2 complexes were only detected in the 200 kd complexes (Fig. 5C, panel 11, fractions 25-27), consistent with previous studies (8, 23). The presence of a putative cyclin D-cdk6 dimer in our system distinguishes cdk6 from cdk4, which appears unable to associate stably with cyclin D in the absence of an assembly factor and is not detected in a dimeric form. We found that cdk6 associated with both cyclin D1 and cyclin D2 in ternary and dimer complexes individually (data not shown), but co-immunoprecipitations are shown for convenience.

We performed cyclin D1/cyclin D2 co-immunoprecipitation analysis with fractions from A, G0 or TGF-β Tet-cdk6^WT^ cells grown in the ON condition (Fig. 5C) and obtained similar results. In each condition, the 200 kd pool contained cdk4, cdk6 and p27 bound to cyclin D1/D2 and the 66 kd pool contained cdk6 bound to cyclin D1/D2 (Fig. 5C, panels 12-22, fractions 25-27 and 30-32, respectively). Cyclin D1/D2 were not associated with cdk4 in the absence of p27 and cyclin D1/D2-cdk4 complexes were not detected in any fractions besides the 200 kD pool in any of the conditions tested (Fig. 5C, panels 12, 16 and 20, fractions 30-32). Additionally, the levels of cyclin D-cdk6-p27 complexes increased when cdk6 was expressed (A, G0 and +TGF-β cells ON) relative to the amount of ternary complex seen in cells grown in the OFF condition, suggesting that the overexpression of cdk6 could sequester p27 efficiently. The putative cyclin D-cdk6 dimer persisted in the presence of TGF-β (Fig. 5C, panel 21, fractions 30-32), suggesting that overexpression of cdk6 exceeded the level of p15 induced by TGF-β treatment.

Next, we analyzed cyclin D1/D2 immunoprecipitates from the parental Mv1Lu cells to verify that the dimer was not an artifact of the tetracycline repressible parent tTA cells (Fig. 5D, panels 23-25). We found that both A and G0 Mv1Lu cells did contain the cyclin D1/D2-cdk6 dimer. Interestingly, TGF-β treatment of Mv1Lu cells caused the loss of both the 200 kD cdk6 ternary complex and the 66 kD putative dimeric complex (Fig. 5D, panel 25). We performed cdk6-associated Rb *in vitro* kinase assays using lysates from A Mv1Lu cells and found that only the ternary cyclin D-cdk 6-p27 complex had catalytic activity (Fig. 5D, panel 26, fraction 26), suggesting that the 66 kD cyclin D1/D2-cdk6 complex was catalytically inactive.

Thus, we had detected additional differences between cdk4 and cdk6: the ability to form a non-cataltyic dimer, which was resistant to TGF-β treatment during cdk6 overexpression conditions. We examined cyclin D1/D2-associated complexes in the cdk6^D163N/R31C^ expressing cells as well. Lysates from A and TGF-β treated cdk6^D163N/R31C^ cells were separated by gel filtration chromatography and the resulting fractions were immunoprecipitated with anti-cyclin D1 and D2 antibodies (Fig. 5E, panels 27 and 28). In contrast to Mv1Lu cells treated with TGF-β, where both the cyclin D-cdk6-p27 ternary and cyclin D1/D2-cdk6 dimeric complexes are dissociated, in the presence of TGF-β in the cdk6^D163N/R31C^ cells both the ternary and the dimeric cdk6 were detected (Fig. 5E, panels 27 and 28, fractions 33 and 34), and the persistence of these pools might account for the ability of the cdk6^D163N/R31C^ cells to continue to proliferate in the presence of TGF-β. The cyclin D-cdk6 dimer was detected in Tet-cdk4^WT^ cells grown in the presence of Tet (cdk4 OFF) (Fig. 5F, panel 29, fractions 29 and 30). However, when the cdk4 protein was expressed (cdk4 ON), the cyclin D-cdk6 dimer was no longer detected (Fig. 5F, panel 30), suggesting that overexpression of cdk4 caused the dissolution of the cyclin D-cdk6 dimer and the redistribution of cyclin D1/D2.

## DISCUSSION

This is the first study that has directly compared the requirement for cdk4 and cdk6 catalytic and sequestration functions in different proliferative conditions, and we have found both functional as well as biochemical differences, which suggest that these two kinases are not functionally redundant as originally presumed. In a kinase-independent manner, non-catalytic cdk6 appears able to sequester p27, activate cdk2 activity, rescue TGF-β-induced growth arrest, and accelerate release from G0, while the homologous cdk4 variant was not competent to perform these actions.

Our data also suggests that once in cycle epithelial cells are able to proliferate in the absence of cyclin D-associated kinase activity, provided that cdk2 kinase activity was maintained, but catalytic cdk4 or cdk6 accelerated release from G0 phase. This suggests that the ability to sequester p27 or other kinase-independent functions may be all that is required of cdk4 and cdk6 during asynchronous proliferation, but a rate-limiting requirement for kinase activity exists upon release from quiescence. Cdk1 or cdk2 might compensate for the loss of cyclin D-associated kinase activity (15, 35), or alternatively, phosphorylation of cyclin D’s catalytic substrates may not be required after their initial activation upon reentry into cycle from G0, or their initial phosphorylation may persist through multiple rounds of division (36, 37). This is consistent with studies using cdk6^−/−^ and cdk4^−/−^ MEFs, which display a reduction in cell cycle reentry following serum deprivation and recent analysis of cdk4/cdk2/Rb triple knockout MEFs that has shown that loss of Rb was able to restore the rate of G0 release to wild type levels (13,14,38). However, we also found that expression of three different non-catalytic alleles of cdk6 was able to increase the rate of G0 exit, suggesting that cdk6 was able to provide a non-catalytic function in this setting as well. It is possible that overexpression of WT cdk6 and the subsequent acceleration of G0 release is not due to increase cyclin D-associated kinase activity, but may be due to the mere presence of the cdk6 protein and its kinase-independent functions.

Additionally, we had originally hypothesized that in the presence of TGF-β, overexpression of either a cdk4 or a cdk6 sink should restore proliferation in the absence of cyclin D kinase activity but this was not observed. Only the expression of sequestration-only cdk6 was able to restore proliferation in the presence of TGF-β, by preventing liberated p27 from associating with cdk2, suggesting that cdk6 was unique. Data suggests that cdk6 expression may provide an additional benefit beyond the maintainence of cdk2 activity. We generated cells that overexpressed cyclin E-cdk2, as we felt that this might also provide an adequate sink for p27, but as seen by others, this did not rescue TGF-β-mediated growth arrest (39, 40), suggesting that expression of cdk6 is unique and provided an additional function.

Our data demonstrates the existence of a previously unidentified cyclin D-cdk6 dimer in proliferating and quiescent cells that while not dependent on cdk6 overexpression can be augmented by increased amounts of cdk6. Cdk6’s association with cyclin D1/D2 in the absence of p27 or other assembly factors distinguishes it from cdk4 and might in part explain the differential activity of cdk4 and cdk6. While we cannot rule out that another small molecule is associated with this complex, the size of the complex is suggestive of a dimer. Mv1Lu cells do not express p15 in the absence of TGF-β (4), so it is unlikely that an Ink4 protein is associated with this complex. We have detected this 66kD cyclin D-cdk6 complex in other cell types, including MCF-10A (human mammary epithelial cells (data not shown). The role of the cyclin D-cdk6 dimer is unclear and obviously will be the focus of future investigation, but we propose that one function may be to act as a reservoir of cyclin D. In the presence of increased p27 levels, the cell could draw from this dimeric form, to increase the catalytically active cyclin D-cdk6-p27 200 kd form. Alternatively, the cyclin D-cdk6 dimer may have additional functions, including the ability to modulate cyclin D transcriptional activity or prevent cyclin D’s aggregation. Cyclin D1 is an oncogene and its inappropriate or constitutive expression increases proliferation, causes DNA rereplication, loss of p53 and a subsequent increase in aneuploidy, demonstrating that its activity must be controlled (41–43). Recent studies have demonstrated that both cyclin D1 and cdk6 appear to associate with transcription factors to modulate their activity on the promoters of cell cycle regulated genes, in a kinase-independent fashion (44, 45). The shuttling of cyclin D from a catalytically inactive dimer to a catalytically active ternary complex might represent an interface between cell cycle activity and transcriptional regulation.

The fact that a cyclin D-cdk6 dimer can exist *in vivo* suggests that inherent structural and stability differences must exist between cdk4 and cdk6. Catalytically active cyclin D-cdk4 complexes cannot be readily reconstituted by mixing recombinant cdk4 and cyclin D, and while co-infection of cdk4 and cyclin D1 in insect cells will yield an active dimer, this complex appears unstable (5) Evidence suggests that the dimeric form does not exist *in vivo* and multiple studies have suggested that all cyclin D-cdk4 complexes contain p27, p21 or another stabilizing molecule (5,8,11). Depletion analysis has shown that while p27-free cdk4 complexes exist in the cell, they are also cyclin D-free and catalytically inactive (23,33,34). The three dimensional structures of cyclin D1-cdk4 and a cyclin D3-cdk4 dimers have been resolved, but in order to achieve this, extensive substitutions between cdk4 and cdk6 had to be made (46, 47). Those authors concluded that in general this complex was inherently unstable in the absence of an assembly factor (46, 47). In contrast, cdk6, with its larger cyclin D interface domain, has been readily crystallized in association with a Kaposi Sarcoma vial cyclin V, p16, p18 and p19 (48–52). Consistent with the idea that cyclin D-cdk4 complexes are unstable compared to cyclin D-cdk6 complexes, purification of catalytically active cyclin D-cdk4 complexes can only be achieved with mild detergents, while cyclin D-cdk6 complexes can be purified under more stringent conditions (53).

In contrast to our findings, other groups have examined the composition of cdk6 complexes in hematopoetic cell lines, where cdk6 is the dominant kinase and/or Ink proteins are constitutively expressed, and in these cases, the cyclin D-cdk6 dimer was not detected (33). We posit that the pool distribution of the cyclin-cdk complexes may be dynamic and dependent upon the intracellular concentration of the different components (Inks, Cip/Kips, cyclin D, cdk4 and cdk6) (Fig. 6). In epithelial cells such as Mv1Lu cells, where neither Inks nor p21 are normally expressed, cdk6 may be required to sequester cyclin D in an inactive pool until needed. This appears to be a cdk6 specific phenomenon, which may have implications in those cancer cells where cdk6 is overexpressed or Ink4 expression is lost.

**Figure 6.**
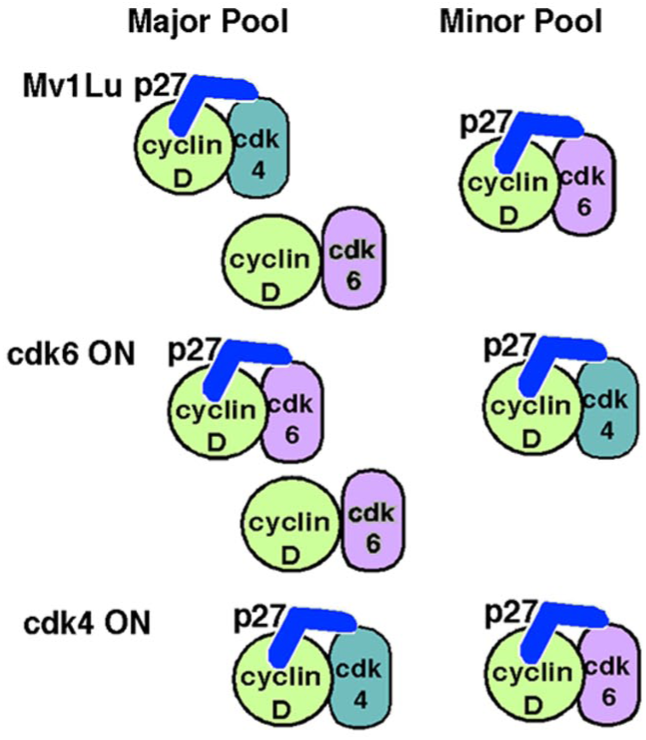
A proposed model of the dynamic nature of cyclin-cdk complexes. Cdk4 and cdk6 exist in multiple complexes that are influenced by the intercellular levels of cyclin D, cdk4, cdk6 and CKIs. In Mv1Lu cells, cyclin D-cdk4-p27 ternary complexes and cyclin D-cdk6 dimers are present as the major pools, while p27-cyclin D-cdk6 ternary complexes are a minor pool. Increasing levels of cdk6 leads to an increase in cyclin D-cdk6-p27 complexes, while increasing cdk4 prevents formation of the cyclin D-cdk6 dimer.

Increased expression of cdk4 blocked formation of the dimer presumably due to increased titration of cyclin D into cyclin D-cdk4-p27 complexes (Fig. 6). When cdk6 was overexpressed, an increase in both the cyclin D-cdk6 dimer and the cyclin D-cdk6-p27 ternary complex was detected (Fig. 6). When cyclin D-cdk6^D163N/R31C^ was overexpressed, which was resistant to p15 induction, it was now able to sequester p27, support cdk2 activity, and maintain the cyclin D-cdk6 dimer. Thus, overexpression of cdk4 or cdk6 altered the overall distribution of the cyclin-cdk complexes, rendering only the cdk6^D163N/R31C^ complexes competent to survive TGF-β-mediated growth arrest. The difference between cdk6’s ability to overcome TGF-β mediated growth arrest may thus be two fold: the reactivation of cdk2 due to increased sequestration by the newly formed cyclin D-cdk6-p27 ternary complex and the persistence of the cyclin D-cdk6 dimer. The fact that overexpression of cyclin E and cdk2 cannot overcome TGF-β mediated growth arrests suggest that the persistence of the dimer may be required for proliferation and maintenance of cdk2 activity alone is not sufficient.

Our study has detected differences between cdk4 and cdk6, consistent with the emerging idea that these two kinases have non-redundant functions. Overexpression of cdk6, but not cdk4, induced astrocyte differentiation, blocked MEL cell differentiation and increased entry of embryonic stem cells into S phase (54–56). Tissue specific defects exist in cdk4 or cdk6 null animals, respectively, despite the existence of the other homologue, arguing for cell type specific functions (13–16). As there are currently small molecule inhibitors that target both cdk4 and cdk6 catalytic activity in cancer clinical trials (57), it will be important to further understand the differences between cdk4 and cdk6, the balance of their kinase dependent and independent roles, and whether inhibiting one function will be an effective enough therapy to halt the growth of cancerous cells.

## ACKNOWLEDGEMENTS

We wish to thank Drs. Sander van den Heuvel and Charles Sherr for the kind gifts of cdk4 and cdk6 plasmids and Dr. Joan Massagué for the kind gift of antibodies. We wish to thank Dina Leznova, Dr. Ting-Chung Suen, Dr. Nuria Ferrandiz-Diaz, Sadia Bhuyan, Jared Macklin and Irfan Azam for technical assistance.

